# Fine-tuned DNA sequence language models for predicting gene expression in admixed population: A comparison with variant-based methods

**DOI:** 10.64898/2026.11.17.767377

**Authors:** Xiaonan Liu, Yu-Jyun Huang, Junwei Lu, Jiecong Lin, Minoli Perera, Kari North, Anne E. Justice, Xihong Lin, Tamar Sofer

**Author notes:** These authors equally contributed to the work (Xihong Lin, Tamar Sofer).

## Abstract

Predicting gene expression from whole genome sequencing data is essential for elucidating regulatory mechanisms and interpreting functional consequences of non-coding variants to complex diseases. While traditional approaches rely on variant-based linear models, DNA sequence-based language models have gained increasing popularity for their ability to learn regulatory features directly from whole genome sequence data. Previous studies have evaluated models such as Enformer, but newer sequence-to-function models, including Borzoi and AlphaGenome, remain less well characterized for individual-level gene expression prediction, particularly in non-European and admixed populations. Here, we evaluated and fine-tuned Borzoi and AlphaGenome using personal genomes paired with whole-blood RNA-seq data from the Hispanic Community Health Study/Study of Latinos (HCHS/SOL). Fine-tuning substantially improved the performance of both sequence-based models compared with their pretrained versions; however, their performances remained similar to that of the variant-based linear models. Leveraging the ancestral diversity of HCHS/SOL, we found that Borzoi and AlphaGenome showed similar performance across Amerindian-, European-, and African-enriched groups. Fine-tuned Borzoi, fine-tuned AlphaGenome and elastic net performed modestly better when trained and evaluated in the same ancestry-enriched group. Finally, we explored strategies to improve prediction performance and found that incorporating predicted epigenetic features improved prediction near the promoter region. Significant progress remains needed before the full potential of sequence-based models is attained. Incorporating more biology-guided training approaches could be helpful to achieving better performance.

## Introduction

Gene expression is an important molecular phenotype linking genetic variation to complex traits. Integrating gene expression with genetic and phenotypic data can help identify genes associated with complex traits through transcriptome-wide association studies (TWAS) ^1,2^ and interpret the functional consequences of noncoding variants through expression quantitative trait locus (eQTL) studies ^3–5^. Current large-scale human genetic studies have extensive multi-ancestry whole-genome sequencing and phenotypic information but generally lack matched transcriptomic data. Therefore, having computational models that can accurately predict individual-level gene expression from whole genome sequence (WGS) could increase the sample size and statistical power of expression-based association analyses and facilitate biological discovery.

Currently, gene expression prediction models fall in two categories (Figure 1), traditional variant-based models and sequence-based models. Traditional models are variant-based, where statistical/machine learning methods are used to regress genetic variant dosages on the expression of a given gene. For example, PrediXcan ^1^ uses elastic net ^6^, FUSION ^2^ evaluates several methods, including best linear unbiased predictor ^7^ and Bayesian sparse linear mixed model ^8^.Although nonlinear approaches such as random forests ^9^ and neural networks ^10^ have been explored, comparative studies have shown that using sparse statistical models generally achieve better variant-based prediction of gene expression ^1,2,11^.

**Figure 1.**
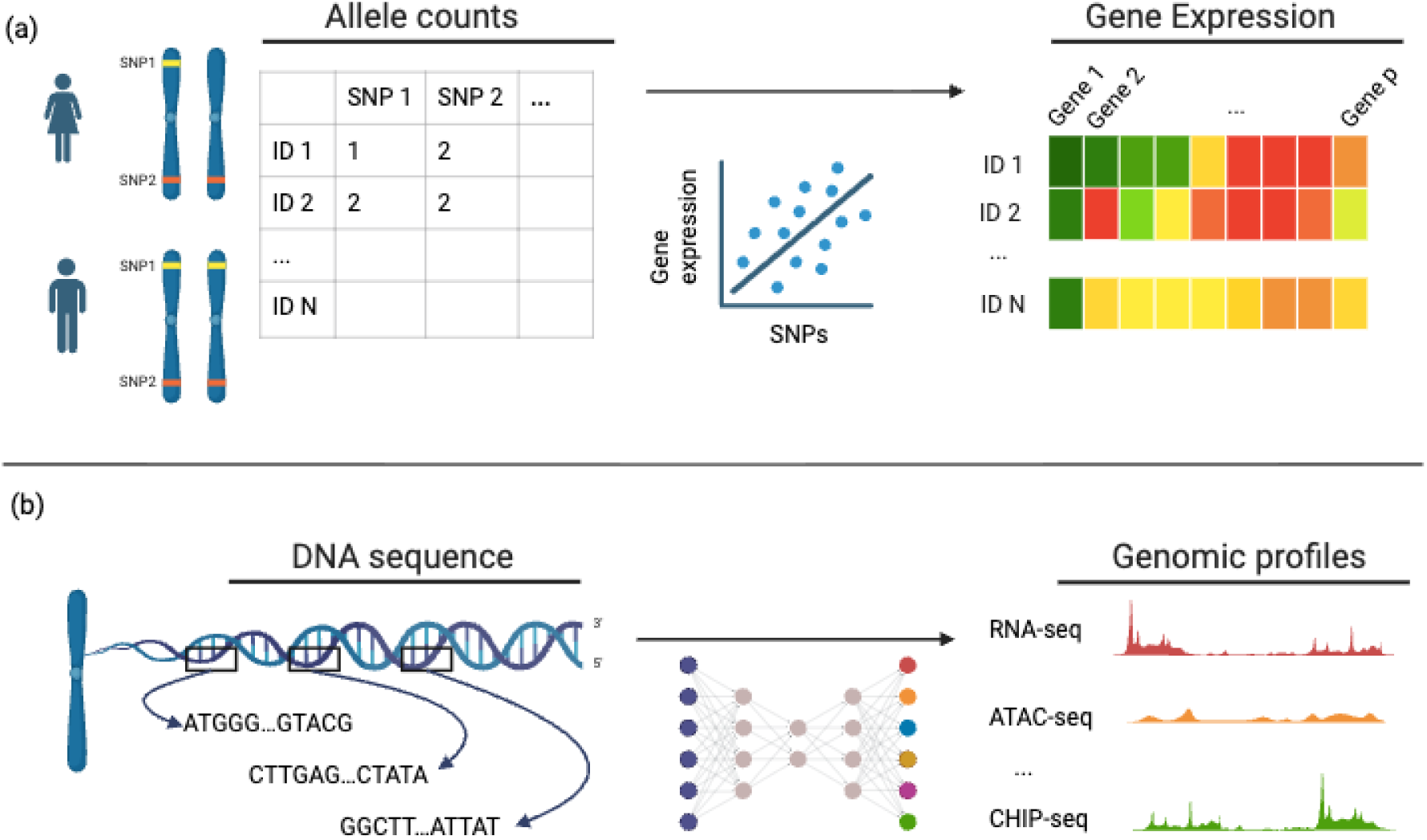
Methods of predicting gene expression: (a) variant based model where linear models are used to regress gene expression on single-nucleotide genetic variants across individuals and prediction is made by aggregating the effects of variant variants using elastic net. (b) sequence-based models where deep learning architectures (e.g. transformers) are used to model phased DNA sequences to predict genomic profiles (e.g. RNA-seq, ATAC-seq, and CHIP-seq-based genomic features).

Recently, inspired by the rapid development of natural language processing, there has been a growing interest in development of sequence-based models that leverage deep learning architecture to learn sequence-to-expression relationships ^12^. These models are trained on reference genome across loci to predict various genomic features, including RNA-seq, CAGE-seq, DNase/ATAC-seq, and CHIP-seq derived genomic features across tissues and cell types. While numerous models exist, recent examples include Enformer ^3^, Borzoi ^5^, and AlphaGenome ^13^.

Multiple groups have benchmarked ^14–16^ and fine-tuned ^17–19^ sequence-based models in predicting individual-level gene expression using eQTL study cohorts, such as GTEx ^20^, ROSMAP ^21^, and GEUVADIS ^22^. Our work contributes to the evaluation of sequence-based gene expression prediction models in several ways. First, previous studies focused primarily on Enformer, we benchmark and fine-tune more recent models, including Borzoi and AlphaGenome. Second, previous analyses were performed primarily in European-ancestry datasets with modest sample size (∼700 individuals), whereas our study leverages a substantially larger cohort of approximately 7,000 individuals from an admixed population. This allows us to assess ancestry-specific differences in model performance and test whether models trained within specific ancestry-enriched groups generalize to admixed population. Third, we investigate strategies to improve predictive performance, providing insight into how these models can be more effectively adapted for individual-level gene expression prediction.

## Methods

### Data

We used individual-level paired WGS and whole blood RNA-seq data from the Hispanic Community Health Study/Study of Latinos (HCHS/SOL) study, with whole genome sequencing performed by the Trans-Omics for Precision Medicine (TOPMed) program ^23^. The HCHS/SOL is a population-based cohort study designed to investigate the chronic disease risk factors among Hispanic/Latino adults in the United States ^24,25^. During 2008-2011, 16,415 participants aged 18-74 years were recruited from four study sites: San Diego, CA; Chicago, IL; Bronx, NY and Miami, FL. At the baseline exam, participants provided blood samples, which were used to measure blood count panels. Whole blood PAXgene RNA tubes were stored at the study’s central laboratory at Minnesota University until they were shipped to the RNA sequencing lab.

#### Whole genome sequencing (WGS)

We obtained the WGS data for 7,729 individuals in HCHS/SOL study from the TOPMed Freeze 8 dataset. Sequencing was performed at the Baylor Human Genome Sequencing Center and aligned to the GRCh38 human reference genome. We excluded, sequentially, 10 duplicated samples, 735 samples without RNA-seq, 241 potential mismatch samples with the RNA-seq (see Supplementary Methods), and 1,623 related individuals. The genetic relationship matrix (GRM) is an N*N matrix of relatedness information of the samples included in the study ad was computed using ‘PC-relate’ R package (version: 1.24.0) ^26^. Unrelated individuals were selected using the sparse GRM by defining relatedness blocks based on nonzero off-diagonal GRM entry and retaining the first individual from each block. The final analysis population comprised of 5,120 independent individuals.

For variant quality control (QC), we excluded variants that: 1) did not pass TOPMed QC filters, 2) were not single nucleotide variants, 3) were monomorphic, 4) had mean sequencing depth 10. To reconstruct individual’s genomes for sequence-based models, we phased WGS using SHAPEIT5 ^27^. To improve phasing accuracy, we augmented our dataset with unrelated TOPMed participants that were not in HCHS/SOL study. We first estimated the distribution of predicted genetic ancestry groups in HCHS/SOL and used these proportions to sample 10,000 TOPMed individuals to approximate the ancestry composition of HCHS/SOL. Genetic ancestry of TOPMed were estimated based on genotype using a semi-supervised machine learning method as described in ^28^.

#### RNA sequencing

Bulk RNA was extracted from whole-blood samples and sequenced at New York Genome Center according to the TOPMed RNA sequencing protocol ^29^. Raw reads underwent QC and were aligned to the human reference genome GRCh38 using STAR (v2.6.1d) ^30^.

Transcript and gene expression quantification were performed using RSEM (v1.3.1) ^31^. Gene quantification and quality control was performed using RNA-SeQC (v2.3.3) ^32^. Expression were generated at both the gene and transcript levels. We selected autosomal genes that were expressed based on the following thresholds: 0.1 Transcripts Per Kilobase Million (TPM) in 20% samples and 6 unnormalized reads in 20% samples. We further retained genes for which the full 1Mb window centered at the transcription start site (TSS) fell within the chromosome, using chromosome lengths obtained from the hg38 FASTA index (.fai) file. Genes whose TSS 1Mb window extended beyond the chromosome length were excluded. Gene and transcript annotations were based on the GTF V34 annotations ^33^.

We normalized gene expression using trimmed mean of M values (TMM) ^34^, consistent with the GTEx pipeline. For model training, tunning, and evaluation, we split the data into training (70%, N=3584), validation (15%, N=768), and test (15%, N=768). TMM normalized expression was regressed on study sites, sex, 11 genotype ancestry principal components, lab-provided proportions of neutrophils, lymphocytes, monocytes, eosinophils, and platelet count. The regression model was fit using the training set only. The fitted coefficients were then applied to training, validation, and test sets, and residual was computed and used as the outcome in downstream genetic prediction models.

### Variant-based model: Elastic net

Variant-based models associate gene expression with genetic variants typically through linear models. This variant-based method offers interpretable results and can model long genomic regions since it focuses on variants instead of raw DNA sequences.

For each gene, we considered local variants within 1Mb region upstream of the TSS and the 1Mb region downstream of the TSS ^35^. We fit an elastic net model ^6^ to identify the variants associated with expression, setting 0.5 to balance between lasso (ℓ_1_) and ridge (ℓ_2_) penalty. The regularization parameter λ was selected based on minimizing the mean-squared-error (MSE) between predicted and observed gene expression in the validation dataset. Analyses were run in R version 4.4.3 (glmnet R package).

### Sequence-based models

Among sequence-based models, we selected Borzoi and AlphaGenome for this analysis because both directly predicts RNA-seq as gene expression. Borzoi takes a 524kb input sequence and outputs 7,611 human genomic tracks, including RNA-seq, CAGE-seq, ATAC/DNase-seq, and CHIP-seq, across the central 196kb region at 32bp resolution. AlphaGenome takes a 1Mb input sequence and outputs 5,930 human genomic tracks, with the additional prediction of PRO-cap, splice sites, splice site usage, splice junctions, DNA contact maps, spanning the full input region. The resolution of AlphaGenome outputs depends on assay type: CHIP-seq is predicted at 128bp resolution, contact maps at 2,048bp, and most other tracks at 1bp resolution ^13^.

Because we use individual genomes stored as variants coded to the reference genome, rather than individual sequences, we implemented a unified computationally-efficient pipeline to derive individual sequences from the reference genome and their genetic variants. For each gene, we extracted DNA sequence of length *L* centered at the TSS ^33^, where *L* =524*kb* for Borzoi and *L* =1 for AlphaGenome. We then applied individuals’ genetic variants to the reference genome by identifying linking variant positions to the reference genome and updating the relevant reference allele to the individual one. Because we used phased data, we reconstructed two sequences for each individual, corresponding to two chromosomal copies. An individual’s DNA sequences is denoted by *x*. We then applied the corresponding model to obtain predictions *ŷ* = *f*(*x*).To derive a gene-level prediction, we selected the output tracks corresponding to RNA-seq in whole blood or K562 cell lines (*T_blood_*). Whole-blood tracks were selected to match the tissue of the observed RNA-seq data, while K562 provides a complementary, more cell-type-specific regulatory context. For each haplotype, we aggregated predictions across output positions overlapping with the exons of the target gene 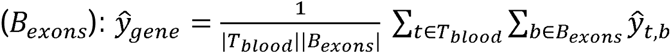 where *b* represents model output positions, corresponding to 32bp bins for Borzoi and individual base pairs for AlphaGenome. Finally, the gene-level predictions from two haplotypes were averaged to obtain each individual’s predicted gene expression. We used gReLU ^36^ python library to run, and fine-tune Borzoi and JAX-based AlphaGenome Research Github codebase to run AlphaGenome, and AlphaGenome PyTorch Github codebase to fine-tune AlphaGenome.

### Fine-tuned Borzoi

Since the original Borzoi was trained using the reference genome, we further fine-tuned the model using individual genomes to improve its performance. We fine-tuned Borzoi separately for each gene with the following changes:

1. **Input sequence length**: Reduced from 524kb to 49,152 bp to enable faster and larger batches within available GPU memory. We removed the cropping layer to accommodate this change.
2. **Output layer**: Modified the last layer to output a single task focused on gene expression prediction.
3. **Pooling function**: Added a pooling function to average predictions around 1kb near the TSS. Borzoi predicts high-resolution genomic tracks without this pooling step. Because our outcome is gene-level expression, we aggregate the local predictions to obtain gene-level predictions.
4. **Diploid inference**: Modified the inference step to compute prediction as the average of predictions from both haplotypes.

We implemented Low Rank Adaptation (LoRA) ^37^ as the fine-tuning method with rank=8 on the attention blocks while keeping the weights of other layers same as the original model (Figure 2). Training hyperparameters were set as follows: learning rate=10^-4^, maximum epochs=50, batch size=20, and gradient accumulation batches=10. To prevent overfitting, we used early stopping with a patience of 5 epochs, terminating training if Spearman’s correlation in validation set did not improve by at least 0.0005.

**Figure 2.**
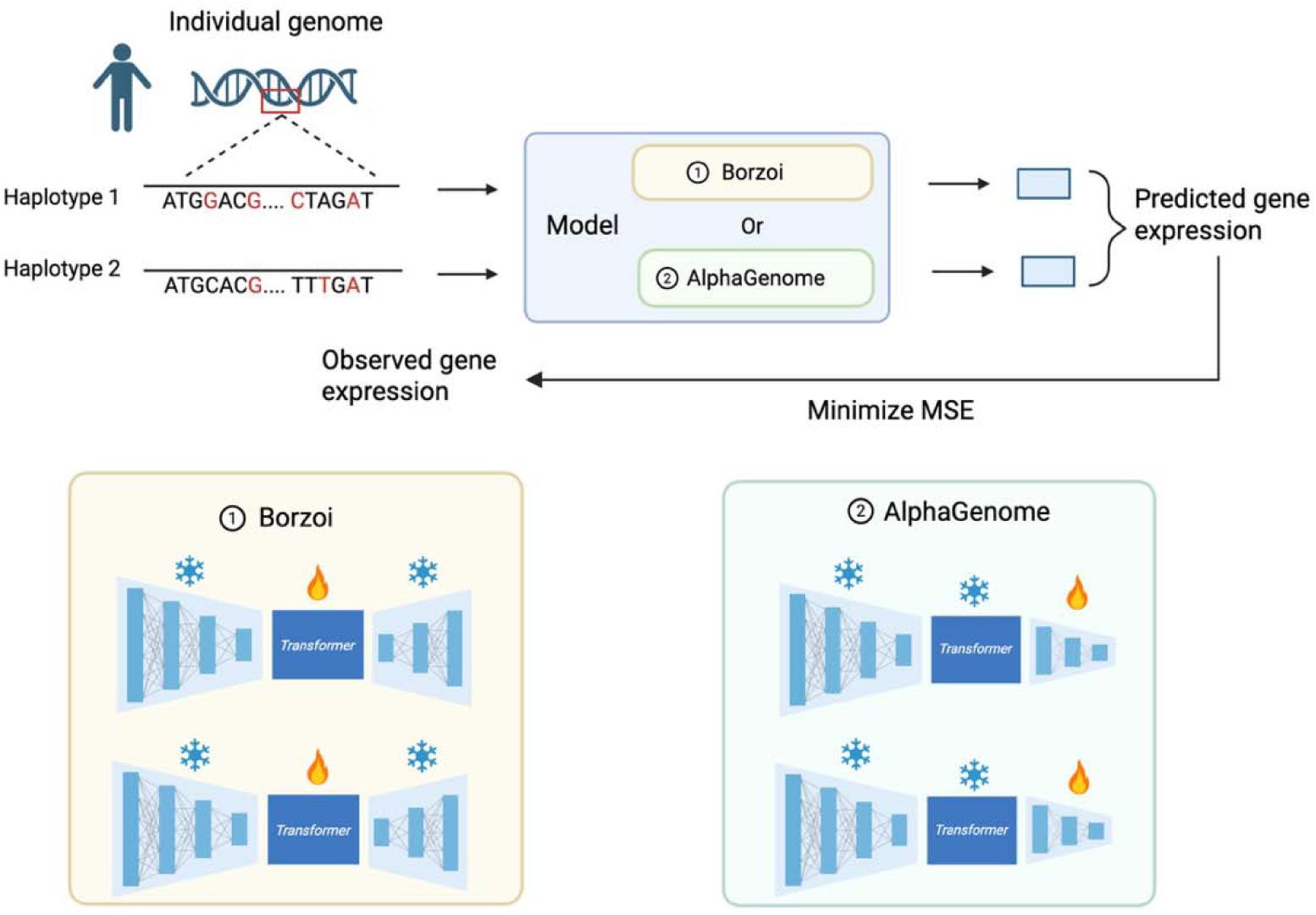
Method for fine-tuning Borzoi and AlphaGenome. The main modifications to Borzoi involve adapting the model architecture to support two input sequences per individual and change the last layer to output a single task focused on gene expression prediction. During fine-tuning, we applied LoRA to the attention blocks while keeping all other parameters same as the original model (i.e. frozen).

### Fine-tuned AlphaGenome

We fine-tuned AlphaGenome using the approach described in ^38^. In details, we used an input sequence length of 49,152bp and extracted the AlphaGenome encoder and transformer layers and added a regression head to predict gene expression. The regression head takes the raw AlphaGenome transformer output, applies normalization followed by dropout (rate=0.05), then uses a linear layer to learn a per-position score. These scores are transformed into weights using a softmax function, which are then used to compute a weighted sum across positions, pooling the features into a scalar prediction.

We fine-tuned AlphaGenome using two approaches: 1) freezing the encoder and transformer layers (i.e. keeping them as they are) and training only the regression head (“head only”), and 2) freezing the pretrained backbone weights while applying LoRA to the transformer attention blocks and training the regression head. Because LoRA did not substantially improve model performance, we proceeded with the head-only fine-tuning approach. Training hyperparameters were set as follows: learning rate=10^-4^, maximum epochs=50, batch size=20, and gradient accumulation batches=10. To prevent overfitting, we used early stopping with a patience of 10 epochs, terminating training if Spearman’s correlation in validation set did not improve by at least 0.001.

### Evaluation

We compared the following models: Borzoi, AlphaGenome, fine-tuned Borzoi, fine-tuned AlphaGenome, and Elastic Net. Model performance was evaluated using Spearman’s correlation between observed and predicted gene expression for each gene, computed on the test dataset. Fine-tuned Borzoi, fine-tuned AlphaGenome, and elastic net were trained on a per-gene basis, with a separate model trained for each gene.

Given the computational constraints of training models for 15,973 genes that passed the minimum expression thresholds, we implemented a strategic sampling approach. We hypothesized that model performance would correlate with *cis*-heritability, which quantifies the proportion of expression variance explained by the additive effects of *cis* genetic variants. To obtain a representative set of genes spanning the performance spectrum while remaining computationally efficient, we selected a subset of 150 genes stratified by *cis*-heritability.

Specifically, we used GCTA ^39^ to compute a *cis*-heritability of each gene based on common variants (minor allele frequency, MAF>0.01) from 1Mb upstream of TSS to 1Mb downstream. We classified genes into three categories: Low (<0.2), medium (0.2-0.5), and high ( 0.5) *cis*-heritability. We then randomly selected 50 genes from each of the categories, yielding 150 genes in total.

### Ancestry-enriched model evaluation

We stratified the study population to “European(EUR)-enriched”, “African(AFR)-enriched”, “Amerindian(AMR)-enriched”, and “Asian(ASN)-enriched” based on proportion of predicted genetic ancestry 50% estimated using the software ADMIXTURE ^40^ (see Supplementary Method). Borzoi and AlphaGenome were applied as pre-trained models without additional training in ancestry-enriched groups. Their prediction performance was evaluated separately within each ancestry-enriched group to determine whether model accuracy differed across groups.

For fine-tuned Borzoi, fine-tuned AlphaGenome and elastic net, we fit separate models within each ancestry-enriched group using only training samples from the corresponding group. To control for differences in sample size across ancestry-enriched groups, we down sampled all groups to the size of the smallest group prior to model training and validation. We primarily assessed how a model trained in one ancestry-enriched group generalizes to the other ancestry-enriched group. We focused on EUR- and AFR-enriched groups because previous studies of polygenic risk score prediction observed large reductions in transferability between EUR-and AFR-ancestry populations ^41^. As a secondary analysis, we compared ancestry-matched performance across EUR-, AMR-, and AFR-enriched groups and evaluated each ancestry-enriched model in the full admixed population (Supplementary Figure 1).

## Result

We computed *cis*-heritability for all 15,973 genes, which showed a broad distribution (mean=0.18, SD=0.14) (Supplementary Figure 2). Based on these estimates, 11,009 genes were classified as low heritability, 4,285 as medium heritability, and 679 as high heritability. From each *cis*-heritability group, we randomly picked 50 genes from each category for model evaluation.

Among these 150 genes, the pretrained sequence-based models showed poor performance in predicting individual-level gene expression in the test set, with low Spearman correlations for both Borzoi (mean Spearman’s *ρ* = 0.11, SD= 0.23) and AlphaGenome (mean *ρ =* 0.15, SD= 0.25). Fine-tuning substantially improved both models, increasing the mean *ρ* to 0.33 (SD=0.26) for Borzoi and 0.29 for AlphaGenome (SD=0.24). Fine-tuned Borzoi outperformed the original Borzoi for 129 genes and exceeded fine-tuned AlphaGenome for 114 genes. However, its performance remained lower than the elastic net model (mean *ρ =* 0.46, SD=0.25) with fine-tuned Borzoi outperforming elastic net for only 21 genes (Figure 3). For elastic net, there were 15 genes without predictions due to no variants selected by the model. Model performance increased with gene-level *cis*-heritability (Figure 3b). This trend was observed across all models, but elastic net showed the strongest performance, particularly among high *cis*-heritability genes (mean *ρ*=0.72), followed by fine-tuned Borzoi (mean =0.53), and fine-tuned AlphaGenome (mean *ρ*=0.48). The same ranking held among genes with lower cis-heritability, although the performance differences were smaller with elastic net had the highest correlation (mean *ρ*=0.15), followed by fine-tuned Borzoi (mean *ρ*=0.11) and fine-tuned AlphaGenome (mean *ρ*=0.09). Figure 3c shows 10 randomly selected genes from each *cis*-heritability group. While all methods have similar performance on the low-heritability genes, differences are apparent in the high-heritability genes.

**Figure 3.**
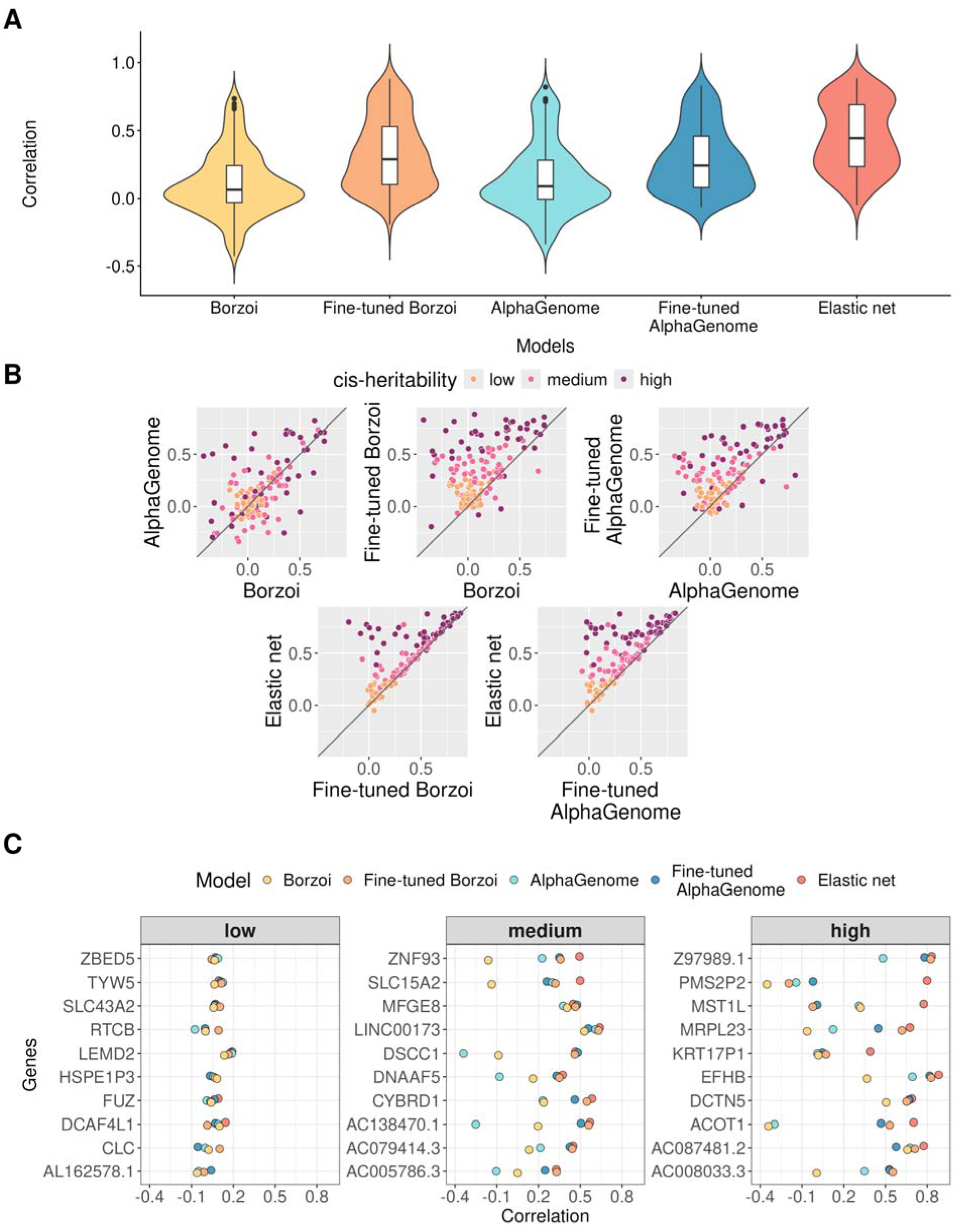
Model performances of Borzoi, AlphaGenome, fine-tuned Borzoi, fine-tuned AlphaGenome, and elastic net. (a) Distribution of Spearman’s correlation across 150 genes evaluated using the test set for Borzoi, AlphaGenome, fine-tuned Borzoi, fine-tuned AlphaGenome and elastic net; (b)Pairwise comparisons of correlation computed using test set by cis-heritability groups. Each dot represents a gene; (c) Correlation of 10 genes randomly selected from low, medium, and high cis-heritability groups respectively.

### Ancestry-enriched model evaluation

Among the study population (N=5,120), there were 3,264 EUR-enriched individuals, 676 AMR-enriched individuals, 469 AFR-enriched individuals, and 4 ASN-enriched individuals. We were unable classify the remaining 707 participants because none of their predicted genetic ancestry proportions exceeded 50%. Given the small sample size for ASN-enriched group, we restricted our analyses to EUR-enriched, AMR-enriched, and AFR-enriched individuals. After down-sampling, there were 305 individuals used for training, 86 for validation, and 78 for testing from each of these groups. Figure 4 shows that Borzoi and AlphaGenome achieved broadly similar prediction performance across three ancestry-enriched groups. Mean Spearman correlations ranged from 0.10 to 0.12 for Borzoi and from 0.13 to 0.15 for AlphaGenome.

**Figure 4.**
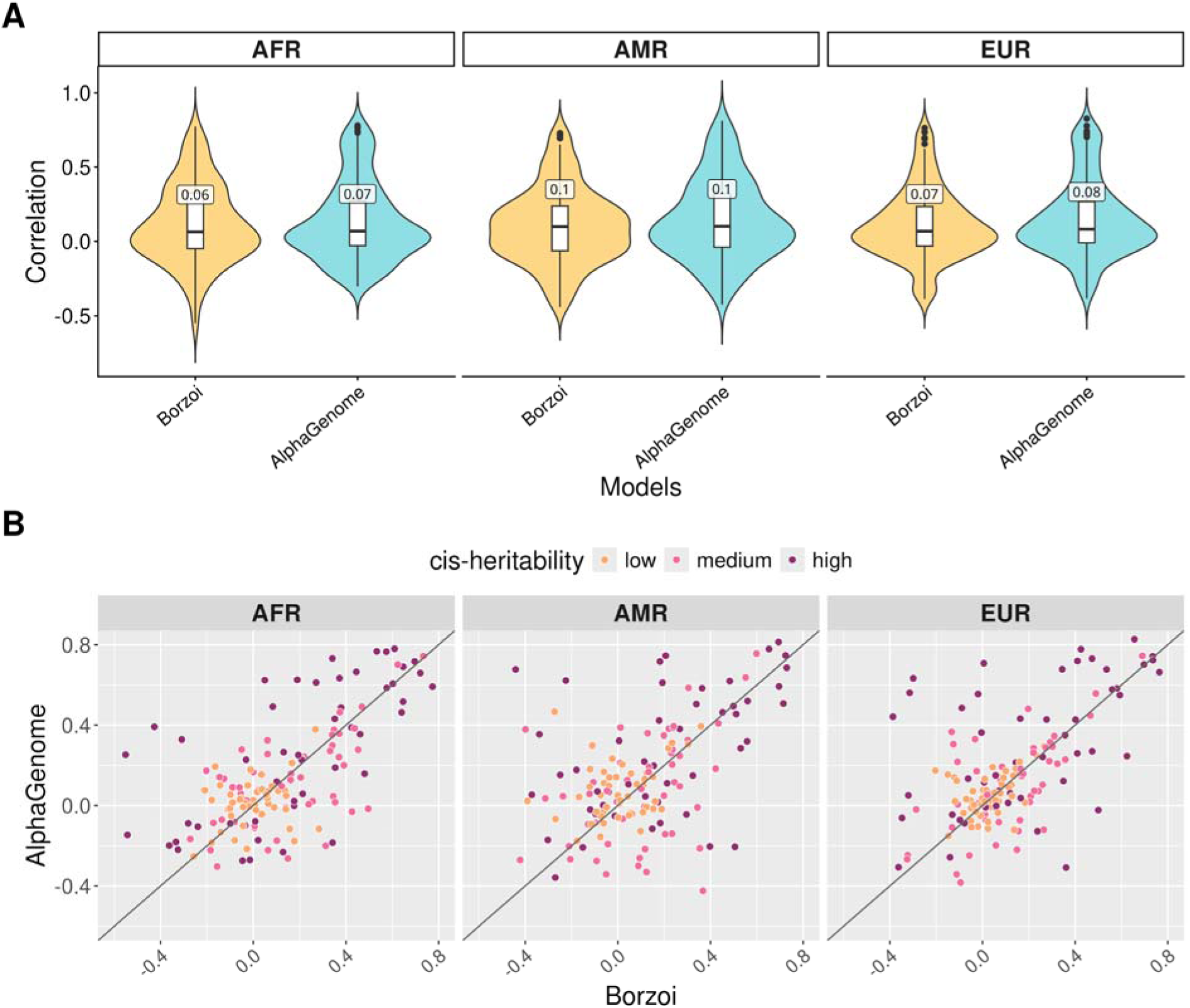
Performance of pre-trained models across ancestry-enriched groups. Both models were pretrained on reference genome only. (a) Spearman’s correlation of Borzoi and AlphaGenome. The median correlation for each model is annotated on the boxplot. (b) Pairwise comparison between Borzoi and AlphaGenome by ancestry-enriched groups and colored by cis-heritability.

Figure 5 compares how models trained in one ancestry-enriched group transfer to other ancestry-enriched groups. Specifically, EUR-trained models were evaluated in EUR- and AFR-enriched individuals respectively, and AFR-trained models were evaluated in both groups respectively. Across all three models, prediction performance was slightly higher when the training and evaluation ancestries were matched.

**Figure 5.**
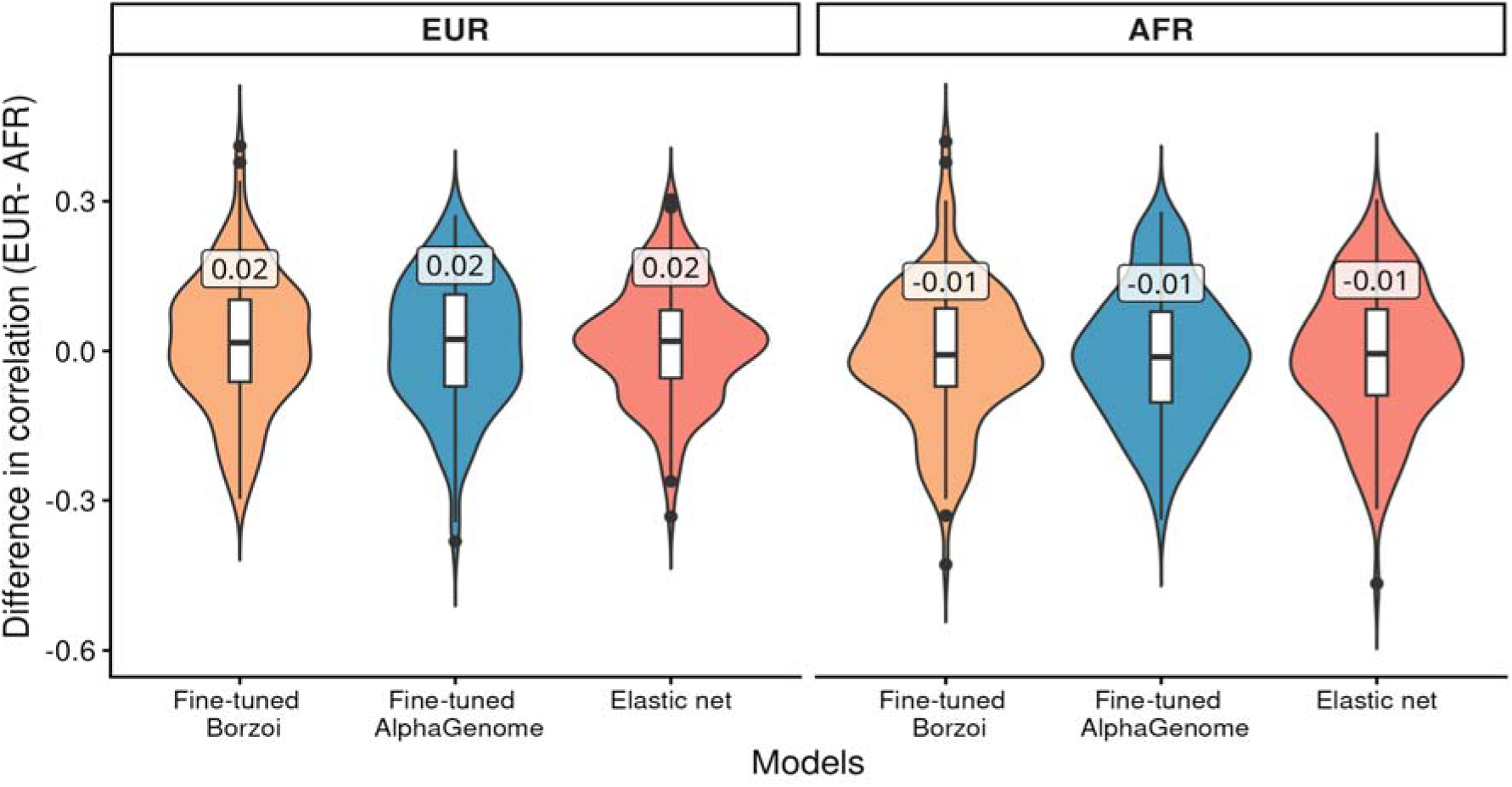
Cross-ancestry differences in performance of ancestry-specific models evaluated in EUR and AFR-enriched groups. Facet labels indicate the ancestry-enriched group used for model training. The y-axis shows the gene-level difference in correlation evaluated on EUR-enriched and AFR-enriched group (EUR-AFR). The median difference for each model is annotated on the corresponding boxplot.

We also evaluated ancestry-matched performance (e.g. training and evaluation in EUR-enriched individuals). Performance was generally consistent across ancestry groups, with elastic net outperforming fine-tuned Borzoi and AlphaGenome (Supplementary Figure 3). Evaluation of ancestry-enriched models in the combined population (e.g. training in EUR-enriched group and evaluation in admixed group) showed similar patterns (Supplementary Figure 4).

### Incorporation of predicted epigenetics

To assess whether gene expression prediction can be further improved, we incorporated predicted epigenetic features from the sequence-based models. We then evaluated whether these features would add information and consequently improve prediction beyond genetic variants alone. We hypothesized that genetic variation explains only part of the gene expression variation, and that epigenetic features such as chromatin accessibility may capture additional regulatory information. We therefore fit elastic net models using three set of features: 1) Single Nucleotide Polymorphisms (SNPs) only, 2) SNPs combined with predicted chromatin accessibility from Borzoi, and 3) SNPs combined with predicted chromatin accessibility from AlphaGenome. Model performance was evaluated across three genomic regions: the promoter region (∼1kb centered at the TSS), 196kb (i.e. output length of Borzoi) and 1Mb (i.e. output length of AlphaGenome) centered at the TSS. For each genomic interval, we extracted the SNPs within the interval and computed the predicted chromatin accessibility over open chromatin regions located in the same interval. Coordinates of open chromatin regions were extracted from ENCODE dataset ENCFF274YGF, which represents open regions derived by DNase-seq in the K562 cell lines.

As illustrated in Figure 6, at the promoter window (∼1kb centered at TSS), adding predicted chromatin accessibility led to clear improvement, with 103 genes showed higher performance with Borzoi-derived accessibility and 84 genes with AlphaGenome-derived accessibility. The mean increases in Spearman correlation is 0.1 for Borzoi and 0.06 for AlphaGenome. However, the benefit of adding predicted chromatin accessibility diminished as the context window increased. At 196kb and 1Mb, performance was similar to that of SNP only model. We further compared gene-expression prediction using SNPs alone with prediction using predicted chromatin accessibility alone (Supplementary Figure 5). At 1kb, Borzoi-predicted accessibility had a higher mean correlation with expression than that predicted using SNPs (0.33 vs. 0.25); at 196 kb and 1 Mb, SNPs performed better than Borzoi- and AlphaGenome-predicted accessibility, respectively (0.43 vs. 0.38; 0.44 vs. 0.34).

**Figure 6.**
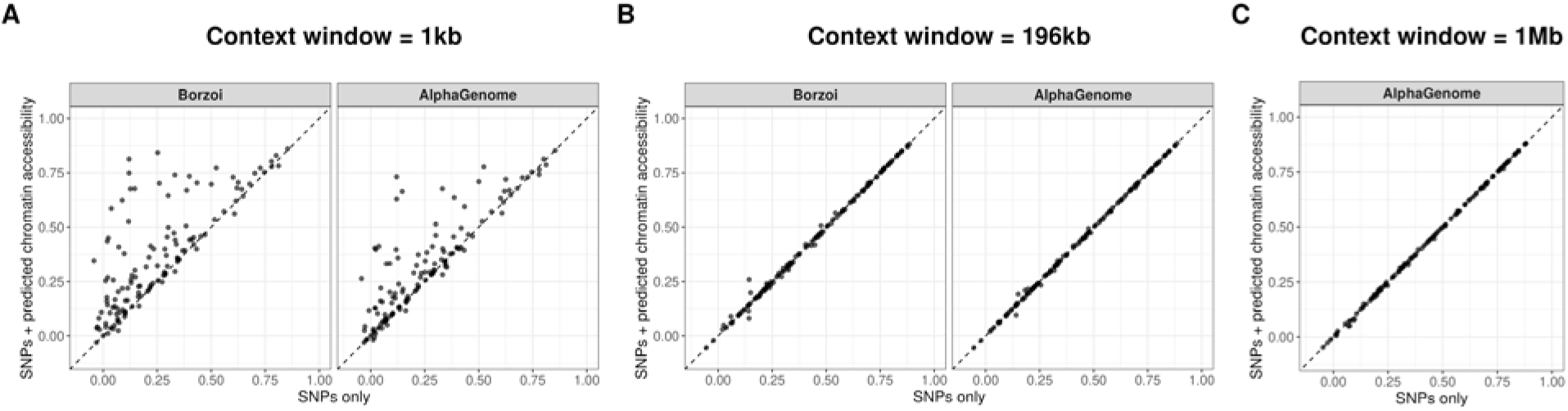
Model performance of elastic net using SNPs only vs SNPs + predicted chromatin accessibility from Borzoi and AlphaGenome. For 1kb, there were 19, 6, 7 genes without predictions for SNPs only, SNPs+pred Borzoi, SNPs+pred AlphaGenome due to no variants selected by the model. For 196kb, there were 11 genes without predictions for SNPs only, SNPs+pred Borzoi, SNPs+pred AlphaGenome due to no variants selected by the model. For 1Mb, there were 12 genes without predictions for SNPs only and SNPs+pred AlphaGenome due to no variants selected by the model.

## Discussion

In summary, we investigated both sequence-based and variant-based models for predicting individual-level gene expression in an admixed population. The sequence-based models, Borzoi and AlphaGenome, showed poor performance in predicting individual-level gene expression. While fine-tuning improved performance, it is comparable to those of linear models. Similar observations have been reported for Enformer by ^14,15,18,19,42^. Adding predicted chromatin accessibility on top of SNPs around TSS improved gene expression prediction but the improvement became less pronounced as context window increased.

From the ancestry-enriched model evaluation, we found training with an ancestry similar to the test set is better at predicting gene expression in test populations. This is consistent with previous evaluations of linear and nonlinear variant-based gene expression prediction models across populations of different ancestries ^43–46^. Rastogi et al. found that fine-tuning Enformer only on European-ancestry individuals performed worse when evaluated in Yoruba-ancestry individuals ^19^. These results highlight the need for expanding RNA-seq data from diverse and admixed populations to improve the multi-ethnic utility of gene expression prediction.

Here we discuss potential reasons why large sequence-based deep learning models, despite their complexity and scale, achieve less than or comparable performance to that of simple linear models. First, these models may not fully capture long-range genomic context. Gene expression is influenced by promoter, distal regulatory elements such as enhancers, and interactions between them. Many sequence-based models were trained using sequences spanning hundreds of kilobases to model distal regulatory interactions.

However, recent studies have shown that while these models capture the effects of promoters, they fail to capture effects of enhancers on gene expression (Karollus, Mauermeier, and Gagneur 2023). In contrast, elastic net directly estimates associations between gene expression and observed genetic variants across the ±1 Mb *cis*-region, allowing it to use distal variants that predict expression without modelling how they exert their effects. Therefore, elastic net’s ability to directly leverage predictive distal variants, together with the limited ability of current sequence-based models to learn distal regulatory effects, may partly explain its stronger performance.

Another possible explanation is that gene expression prediction may benefit from better incorporation of epigenetic information. DNA sequence alone can only explain gene expression to a limited extent. Many other factors can substantially impact gene expression, including chromatin accessibility, 3D DNA contacts, and transcription factor binding. Current sequence-based models use DNA sequence as the sole input and learn to predict multiple genomic profiles, but these intermediate regulatory features may not be fully leveraged for gene expression prediction. Recent studies have shown that explicitly incorporating epigenomic profiles can improve performance. For example, Lin et al. integrated multimodal epigenomic data, including H3K27ac, DNase, and HiC data, with sequence-based models and reported improved gene expression prediction compared to Enformer ^47^. Fu et al. developed a foundation model of transcription that was first pretrained on chromatin accessibility data and subsequently fine-tuned for gene expression prediction, achieving high accuracy in previously unseen cell types ^48^. Because individual-level epigenomic measurements were not available in the TOPMed study, we instead evaluated whether incorporating sequence-model-predicted epigenetic features could improve prediction beyond genetic variants alone. We observed that the performance improved only when using genetic variants and epigenetic features limited to the promoter regions. This may suggest that either the model did not accurately predict chromatin accessibility in distal regions, or that the predicted epigenetic features provided limited information beyond that already captured by the genetic variants.

Our study has several strengths. We evaluated recent sequence-to-function models, including Borzoi and AlphaGenome, extending prior work that has been largely focused on earlier models such as Enformer. We also analyzed gene expression prediction in an admixed population, allowing us to assess model performance cross populations. In addition, we performed analyses to identify potential strategies for improving individual-level gene expression prediction. Our study also has several limitations. First, we used bulk-level, whole-blood, gene expression data. Because gene expression is cell-type specific, bulk measurements average signals across heterogeneous cell populations, making it difficult for models to learn cell-type specific regulatory mechanisms. This limitation may partly explain the modest prediction performance observed. Second, for computational efficiency, we analyzed 150 genes rather than all genes across the genome. Although a larger gene set would provide a more comprehensive assessment, we strategically selected genes to enable evaluating model performance across varying levels of *cis*-heritability. Finally, when constructing individual genomes, we considered only single nucleotide variants and centered model input sequence on the TSS. Future work should investigate whether incorporating indels and structural variants and alternative input generation strategies (e.g. shifted sequence windows) improve prediction.

In conclusion, sequence-based models have emerged as useful approaches for unraveling the regulatory grammar of the genome. Unlike variant-based models, they have the potential to predict effects of unseen variant and learn higher-order interactions among distal regulatory elements. However, fine-tuning two popular sequence-based models using individual genomes improved gene expression prediction but were comparable to linear model performance. Further work is needed before sequence-based models can utilize their full potential and incorporating more biology-guided training approaches could be helpful to achieving better performance.

## Acknowledgement

This work was supported by the NIH grants: R35-CA197449, R01-HL163560, and U01-HG012064, and R01AG080598.

## Reference

1. Gamazon, E. R. et al. (TWAS) A gene-based association method for mapping traits using reference transcriptome data. Nat. Genet. 47, 1091–1098 (2015).

2. Gusev, A. et al. Integrative approaches for large-scale transcriptome-wide association studies. Nat. Genet. 48, 245–252 (2016).

3. Avsec, Ž., et al. (Enformer) Effective gene expression prediction from sequence by integrating long-range interactions. Nat. Methods 18, 1196–1203 (2021).

4. Lal, A. et al. Decoding sequence determinants of gene expression in diverse cellular and disease states. Nat. Methods 23, 1138–1151 (2026).

5. Linder, J., Srivastava, D., Yuan, H., Agarwal, V. & Kelley, D. R. Predicting RNA-seq coverage from DNA sequence as a unifying model of gene regulation. Nat. Genet. 57, 949–961 (2025).

6. Zou, H. & Hastie, T. Regularization and variable selection via the elastic net. J. R. Stat. Soc. Series B Stat. Methodol. 67, 301–320 (2005).

7. Robinson, G. K. That BLUP is a Good Thing: The Estimation of Random Effects. SSO Schweiz. Monatsschr. Zahnheilkd. 6, 15–32 (1991).

8. Zhou, X., Carbonetto, P. & Stephens, M. Polygenic modeling with bayesian sparse linear mixed models. PLoS Genet. 9, e1003264 (2013).

9. Breiman, L. Random forests. Mach. Learn. 45, 5–32 (2001).

10. Ramprasad, P., Ren, J. & Pan, W. Enhancing gene expression predictions using deep learning and functional annotations. Genet. Epidemiol. https://doi.org/10.1002/gepi.22595 (2024) doi:10.1002/gepi.22595.

11. Fryett, J. J., Morris, A. P. & Cordell, H. J. Investigation of prediction accuracy and the impact of sample size, ancestry, and tissue in transcriptome-wide association studies. Genet. Epidemiol. 44, 425–441 (2020).

12. Barbadilla-Martínez, L., Klaassen, N., van Steensel, B. & de Ridder, J. Predicting gene expression from DNA sequence using deep learning models. Nat. Rev. Genet. 26, 666–680 (2025).

13. Avsec, Ž., et al. Advancing regulatory variant effect prediction with AlphaGenome. Nature 649, 1206–1218 (2026).

14. Sasse, A. et al. Benchmarking of deep neural networks for predicting personal gene expression from DNA sequence highlights shortcomings. Nat. Genet. 55, 2060–2064 (2023).

15. Huang, C. et al. Personal transcriptome variation is poorly explained by current genomic deep learning models. Nat. Genet. 55, 2056–2059 (2023).

16. Li, S., Luo, R. & Huang, Y. xDecoder unlocks the potential of genomic foundation models for few-shot personal gene expression prediction. Mol. Syst. Biol. 1–16 (2026).

17. Spiro, A. E. et al. A scalable approach to investigating sequence-to-function predictions from personal genomes. Nat. Methods 1–5 (2026).

18. Drusinsky, S., Whalen, S. & Pollard, K. S. Deep-learning prediction of gene expression from personal genomes. Genome Biol. 27, 19 (2026).

19. Rastogi, R., Reddy, A. J., Chung, R. & Ioannidis, N. M. Fine-tuning sequence-to-expression models on personal genome and transcriptome data. Genome Biol. 1–21 (2026).

20. GTEx Consortium. The Genotype-Tissue Expression (GTEx) project. Nat. Genet. 45, 580–585 (2013).

21. Bennett, D. A. et al. Religious Orders Study and Rush Memory and Aging Project. J. Alzheimers. Dis. 64, S161–S189 (2018).

22. Lappalainen, T. et al. Transcriptome and genome sequencing uncovers functional variation in humans. Nature 501, 506–511 (2013).

23. Taliun, D. et al. Sequencing of 53,831 diverse genomes from the NHLBI TOPMed Program. Nature 590, 290–299 (2021).

24. Sorlie, P. D. et al. Design and implementation of the Hispanic Community Health Study/Study of Latinos. Ann. Epidemiol. 20, 629–641 (2010).

25. Lavange, L. M. et al. Sample design and cohort selection in the Hispanic Community Health Study/Study of Latinos. Ann. Epidemiol. 20, 642–649 (2010).

26. Conomos, M. P., Reiner, A. P., Weir, B. S. & Thornton, T. A. Model-free estimation of recent genetic relatedness. Am. J. Hum. Genet. 98, 127–148 (2016).

27. Hofmeister, R. J., Ribeiro, D. M., Rubinacci, S. & Delaneau, O. Accurate rare variant phasing of whole-genome and whole-exome sequencing data in the UK Biobank. Nat. Genet. 55, 1243–1249 (2023).

28. Shyr, D. et al. Semi-supervised machine learning method for predicting homogeneous ancestry groups to assess Hardy-Weinberg equilibrium in diverse whole-genome sequencing studies. Am. J. Hum. Genet. 111, 2129–2138 (2024).

29. TOPMed_RNAseq_pipeline_COREyr3. Preprint at https://topmed.nhlbi.nih.gov/sites/default/files/TOPMed_RNAseq_pipeline_COREyr3.pdf.

30. Dobin, A. et al. STAR: ultrafast universal RNA-seq aligner. Bioinformatics 29, 15–21 (2013).

31. Li, B. & Dewey, C. N. RSEM: accurate transcript quantification from RNA-Seq data with or without a reference genome. BMC Bioinformatics 12, 323 (2011).

32. Graubert, A., Aguet, F., Ravi, A., Ardlie, K. G. & Getz, G. RNA-SeQC 2: efficient RNA-seq quality control and quantification for large cohorts. Bioinformatics 37, 3048–3050 (2021).

33. Mudge, J. M. et al. GENCODE 2025: reference gene annotation for human and mouse. Nucleic Acids Res. 53, D966–D975 (2025).

34. Robinson, M. D. & Oshlack, A. A scaling normalization method for differential expression analysis of RNA-seq data. Genome Biol. 11, R25 (2010).

35. Araujo, D. S. et al. Multivariate adaptive shrinkage improves cross-population transcriptome prediction and association studies in underrepresented populations. HGG Adv. 4, 100216 (2023).

36. Lal, A., Gunsalus, L., Nair, S., Biancalani, T. & Eraslan, G. gReLU: A comprehensive framework for DNA sequence modeling and design. bioRxiv 2024.09.18.613778 (2024) doi:10.1101/2024.09.18.613778.

37. Hu, E. J., et al. LoRA: Low-Rank Adaptation of large language models. *arXiv [cs.CL]* (2021).

38. Murphy, A., Durán, A. & Koo, P. Adapting AlphaGenome to MPRA data. Genomics × AI Blog https://doi.org/10.5281/ZENODO.20272900 (2026) doi:10.5281/ZENODO.20272900.

39. Yang, J., Lee, S. H., Goddard, M. E. & Visscher, P. M. GCTA: a tool for genome-wide complex trait analysis. Am. J. Hum. Genet. 88, 76–82 (2011).

40. Alexander, D. H., Novembre, J. & Lange, K. Fast model-based estimation of ancestry in unrelated individuals. Genome Res. 19, 1655–1664 (2009).

41. Martin, A. R. et al. Clinical use of current polygenic risk scores may exacerbate health disparities. Nat. Genet. 51, 584–591 (2019).

42. Ramprasad, P., Pai, N. & Pan, W. Enhancing personalized gene expression prediction from DNA sequences using genomic foundation models. HGG Adv. 5, 100347 (2024).

43. Keys, K. L. et al. On the cross-population generalizability of gene expression prediction models. PLoS Genet. 16, e1008927 (2020).

44. Pagnuco, I., Eyre, S., Rattray, M. & Morris, A. P. Transferability of single- and cross-tissue transcriptome imputation models across ancestry groups. Genet. Epidemiol. 49, e22611 (2025).

45. Mogil, L. S. et al. Genetic architecture of gene expression traits across diverse populations. PLoS Genet. 14, e1007586 (2018).

46. Okoro, P. C. et al. Transcriptome prediction performance across machine learning models and diverse ancestries. HGG Adv. 2, 100019 (2021).

47. Lin, J., Li, Z., Zhao, Y., Luo, R. & Pinello, L. EPInformer: scalable and integrative prediction of gene expression from promoter-enhancer sequences with multimodal epigenomic profiles. Nat. Commun. 17, 3975 (2026).

48. Fu, X. et al. A foundation model of transcription across human cell types. Nature 1–9 (2025).

