## Supplementary material for "Fine-tuned DNA sequence language models for predicting gene expression in admixed population: A comparison with variant-based methods"

### Method

#### Mismatch samples

To identify potential genotype-RNA-seq sample mismatches, we first selected the 100 genes with the highest cis-heritability. For each gene, we fit an elastic net model using local variants within 1Mb region upstream of the TSS and the 1Mb region downstream of the TSS. The outcome was inverse-Normal transformed TMM. Sex, study sites, and the first 11 genotype principal components were included as unpenalized covariates. We used 10-fold cross-validation to select the optimal penalty parameter and then generated predicted expression values for each individual for each gene.

As an initial screen for potential sex mismatches, we performed principal component analysis of RNA-seq read counts and visualized the sampled by reported sex. We then compared observed and predicted gene expression across the 100 high cis-heritability genes. For each gene, we plotted the observed and predicted expression and highlighted samples flagged as potential sex mismatches.

To identify additional potential mismatches, we calculate two sample-level metrics across the 100 genes: 1) the root mean squared error (RMSE) between observed and predicted expression, 2) correlation between observed and predicted expression across genes. Samples with poor concordance between predicted and observed expression were flagged as potential mismatches. In details, we used the sex-mismatch samples to define empirical thresholds: the minimum RMSE and maximum correlation among these samples. Individuals with RMSE greater than or equal to this threshold, or correlation less than or equal to this threshold, were classified as potential mismatches.

#### Proportion of predicted genetic ancestry

HCHS/SOL individuals are admixed with, primarily, three ancestral populations: European, African, and Amerindian. A small number of participants (less than 20) have also previously been shown to have substantial levels of Asian ancestry (Conomos et al. 2016). We estimated global ancestry proportions (i.e., proportions of genome inherited from European, African, Amerindian, and Asian ancestry for each individual) using ADMIXTURE (Alexander, Novembre, and Lange 2009).To support the estimation of Asian ancestry and minimize potential miscalibration caused by the small number of HCHS/SOL participants with substantial Asian ancestry, we incorporated whole-genome sequencing data from 1,473 Asian reference individuals from the 1000 Genomes Project (1000 Genomes Project Consortium et al. 2010), the Human Genome Diversity Project (HGDP) (Cavalli-Sforza 2005), and the Simons Genome Diversity Project (SGDP) (Mallick et al. 2016).

Since ADMIXTURE requires unrelated individuals, we divided the data into multiple overlapping subsets, each consisting only of unrelated individuals. Every HCHS/SOL participant with whole-genome sequencing data was included in at least one subset, and ADMIXTURE was applied separately to each subset.

### Figures

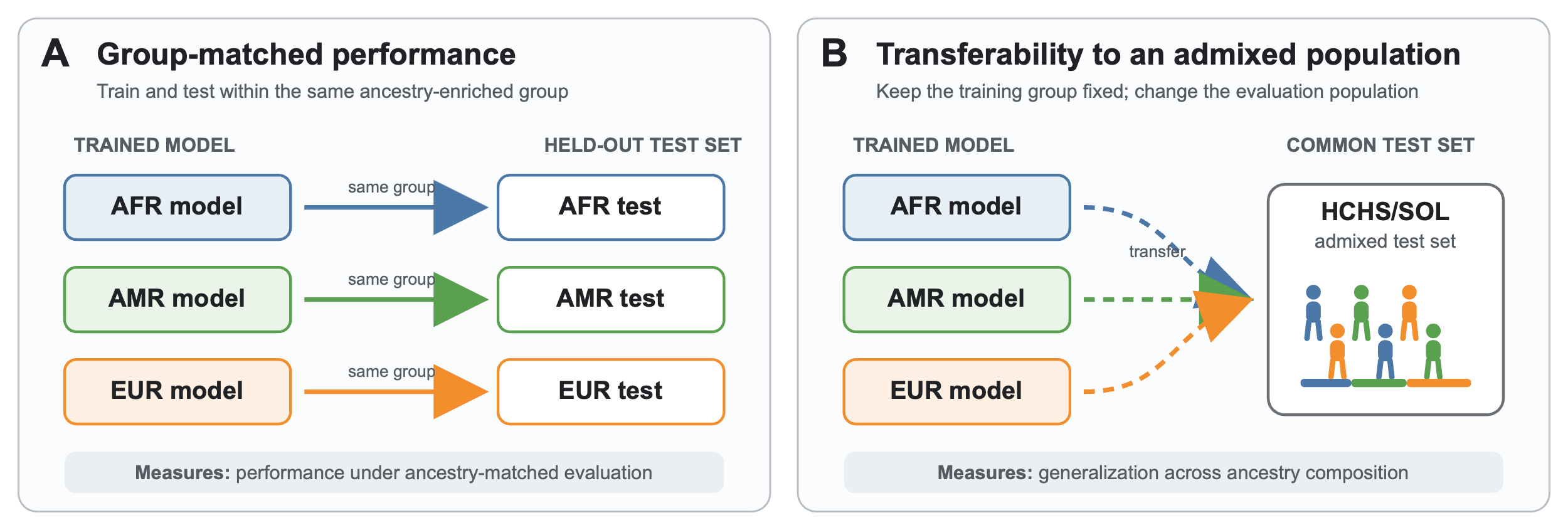

Supplementary Figure 1. Overview of secondary analyses of ancestry-specific model evaluation. We fit separate models within each ancestry-enriched group using only training samples from the corresponding group. We then evaluated each model in two settings: A) on test samples from the same ancestry-enriched group to estimate group-matched performance; B) on test samples from the admixed population to quantify how well ancestry-enriched models generalize across ancestry-enriched groups.

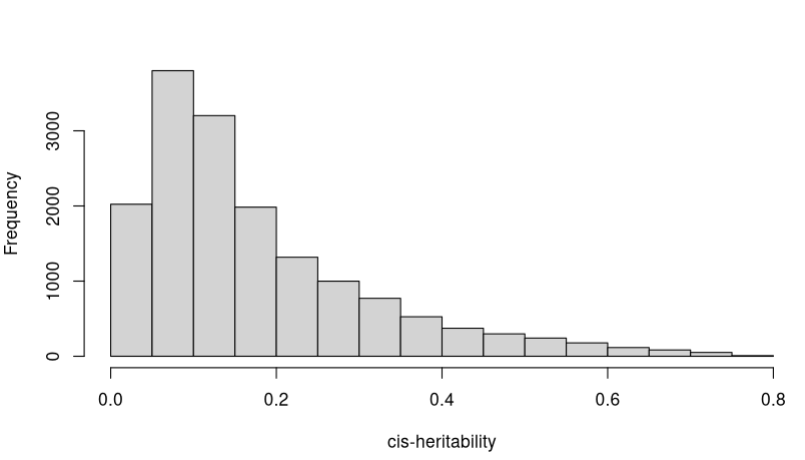

Supplementary Figure 2. Distribution of cis-heritability of 15,973 genes.

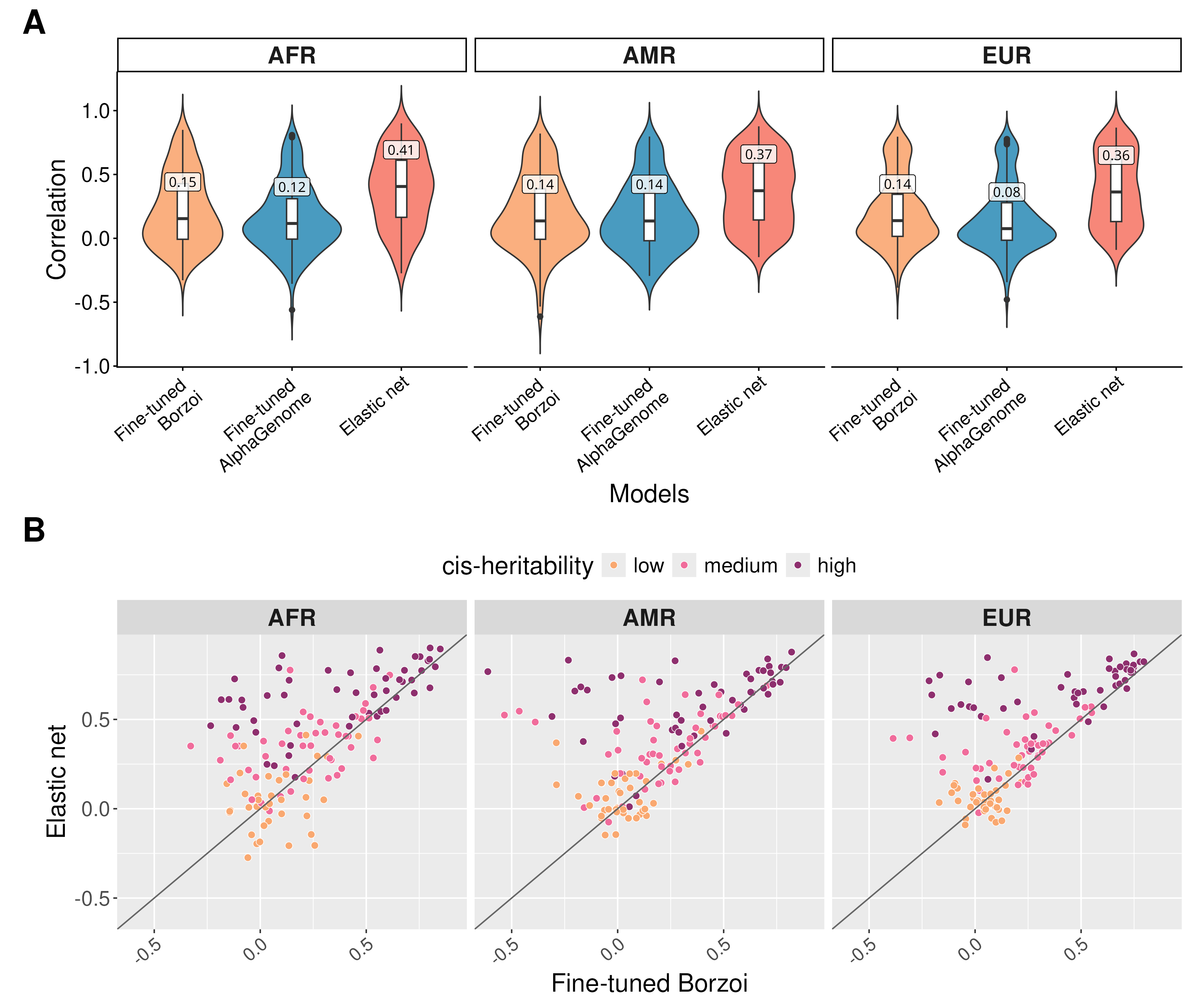

Supplementary Figure 3. Group-matched performance of ancestry-specific trained models. Models trained on AFR-enriched, AMR-enriched, and EUR-enriched groups and evaluated on the corresponding group that were trained on. (a) Spearman’s correlation of fine-tuned Borzoi, fine-tuned AlphaGenome, and elastic net. The median correlation for each model is annotated on the boxplot. (b) Pairwise comparison between fine-tuned Borzoi and elastic net by ancestry-enriched groups and colored by cis-heritability. For elastic net, there were 15,12,17 genes without predictions in AFR, AMR, EUR-enriched groups respectively due to no variants selected by the model.

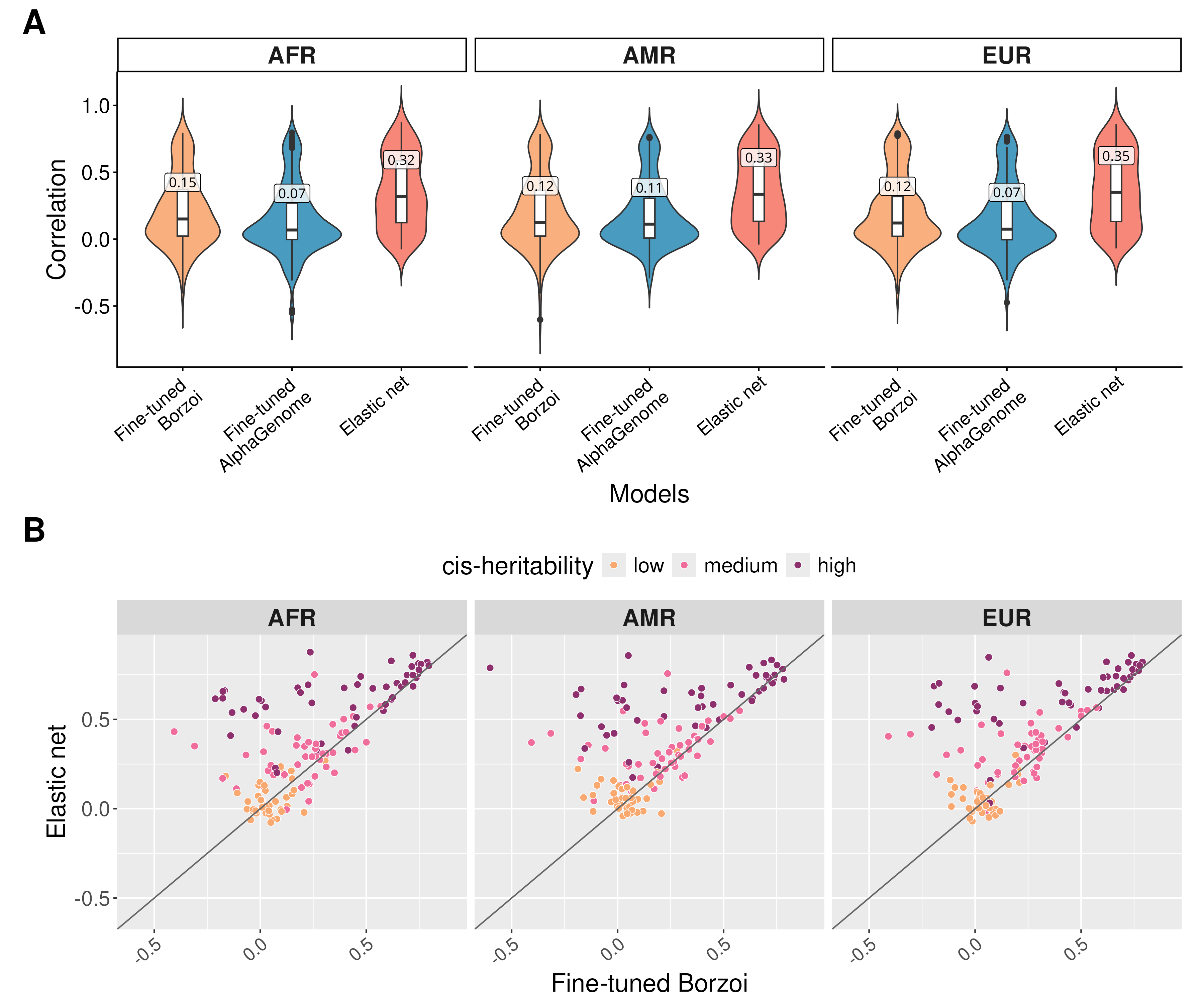

Supplementary Figure 4. Ancestry-enriched models evaluated on entire HCHS/SOL test data. Models trained on AFR-enriched, AMR-enriched, and EUR-enriched groups and evaluated on admixed population. (a) Spearman’s correlation of fine-tuned Borzoi, fine-tuned AlphaGenome, and elastic net. The median correlation for each model is annotated on the boxplot. (b) Pairwise comparison between fine-tuned Borzoi and elastic net by ancestry-enriched groups and colored by cis-heritability. There were 10 genes without predictions from elastic net in AFR-enriched group, 12 in AMR enriched, and 16 in EUR enriched groups due to no associated variants detected by the model.

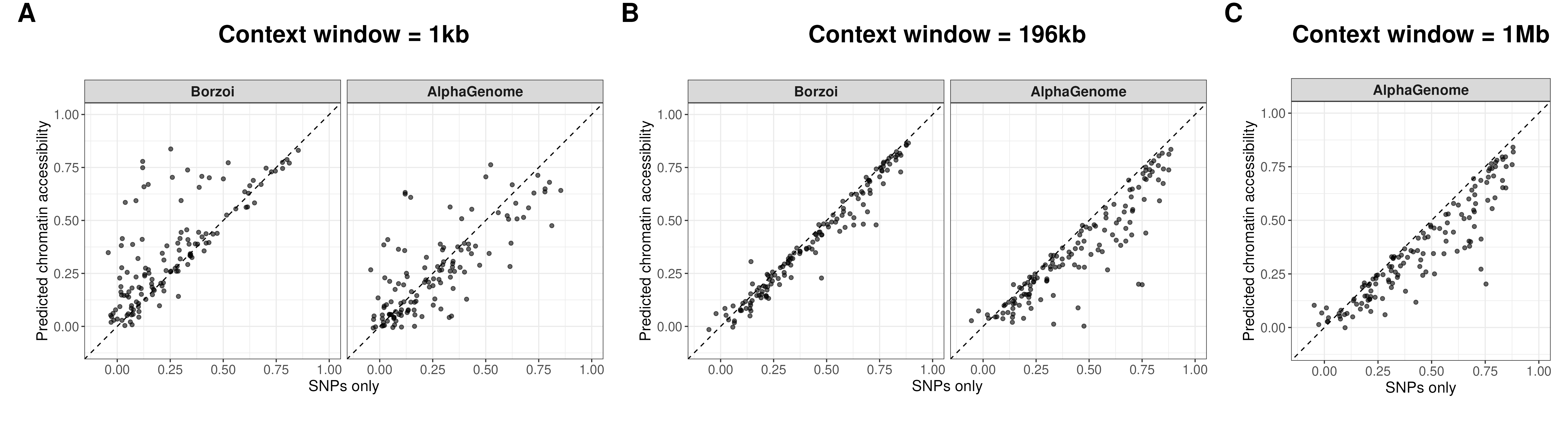

Supplementary Figure 5. Model performance of elastic net using SNPs only vs predicted chromatin accessibility from Borzoi and AlphaGenome.

### Reference

1000 Genomes Project Consortium, Gonçalo R. Abecasis, David Altshuler, Adam Auton, Lisa D. Brooks, Richard M. Durbin, Richard A. Gibbs, Matt E. Hurles, and Gil A. McVean. 2010. “A Map of Human Genome Variation from Population-Scale Sequencing.” *Nature* 467 (7319): 1061–73.

Alexander, David H., John Novembre, and Kenneth Lange. 2009. “Fast Model-Based Estimation of Ancestry in Unrelated Individuals.” *Genome Research* 19 (9): 1655–64.

Cavalli-Sforza, L. Luca. 2005. “The Human Genome Diversity Project: Past, Present and Future.” *Nature Reviews. Genetics* 6 (4): 333–40.

Conomos, Matthew P., Cecelia A. Laurie, Adrienne M. Stilp, Stephanie M. Gogarten, Caitlin P. McHugh, Sarah C. Nelson, Tamar Sofer, et al. 2016. “Genetic Diversity and Association Studies in US Hispanic/Latino Populations: Applications in the Hispanic Community Health Study/Study of Latinos.” *American Journal of Human Genetics* 98 (1): 165–84.

Mallick, Swapan, Heng Li, Mark Lipson, Iain Mathieson, Melissa Gymrek, Fernando Racimo, Mengyao Zhao, et al. 2016. “The Simons Genome Diversity Project: 300 Genomes from 142 Diverse Populations.” *Nature* 538 (7624): 201–6.
